# Impact of winter enclosures on energy expenditure and movement activity of european reed deer

**DOI:** 10.1101/2024.10.19.619248

**Authors:** Václav Silovský, Anna Pilská, Monika Faltusová, Miloš Ježek

## Abstract

The management of red deer enclosures, combined with supplementary feeding, can reduce the energy expenditure of confined animals. These enclosed individuals minimize their spatio-temporal movement and energy usage during their time inside the winter enclosure compared to those living in free range during the same period. The datasets used in this study were based on biologging technology. All measured animals (n=11) were equipped with GPS collars and Daily Dairy sensors to record Vectorial Dynamic Body Acceleration (VeDBA). The study revealed differences in the daily summary and daily mean of VeDBA parameters between enclosed and free-living animals in the study area. Energy expenditure was similar for both groups only at the moment of winter enclosure release. After the enclosure was opened, the released individuals increased their movement and energy expenditure. The use of winter enclosures for red deer management can reduce animal movement and minimize potential damage to young forest stands and seedlings. The opening of the winter enclosure must be synchronized with the availability of natural food resources in the locality. The management of winter enclosures should be carried out with a system of supplementary feeding inside.

## Introduction

In recent decades, Europe has experienced an increase in both the abundance and geographical range of ungulate game populations (Carpio et al., 2021; Heurich et al., 2015; Valente et al., 2020; van Beeck Calkoen et al., 2023). This growth is attributed to climate and environmental changes, and the limited presence of large carnivores has left ungulate populations primarily regulated by food availability and hunting (Heurich et al., 2015; Kuijper et al., 2013; Martin et al., 2020). Hunting has become a major method for managing these populations, but its effectiveness varies based on effort, legislation, and hunting philosophies (Bischof et al., 2012; Heurich et al., 2015; Jarnemo & Wikenros, 2014; Linnell et al., 2020; Möst et al., 2015). The management of deer populations in European forests presents a complex challenge, balancing ecological integrity with conservation and human interests. This usually relates to the state of the environment and especially to the economic damages caused to forests. Their management then includes many activities, which usually relate to supplementary feeding or hunting. In connection with red deer, so-called winter enclosures are often used. In Europe, this is a common practice based on enclosing wild animals in fenced areas of relatively small size (units, max. lower tens of hectares), where the animals are dependent only on food provided by humans. Enclosures are usually closed from the beginning of January to the end of April, i.e., during the period of the most frequent forest damage. This activity likely significantly affects their spatial and behavioral ecology by determining resource allocation and movement barriers. In protected areas, they prevent forest damage and support biodiversity conservation during critical seasons (Riesch et al., 2019). In hunting areas, the management supports population productivity, winter survival, density, and trophy quality (Carpio et al., 2021; Putman et al., 2004). One key aspect of this challenge is understanding the energy requirements and spatial activity patterns of deer across different seasons (Apollonio et al., 2017).

European red deer (Cervus elaphus), especially in winter and spring, exhibit a variety of spatial activities and energy requirements due to changes in environmental conditions and resource availability (Berger et al., 2002; Luccarini et al., 2006; Náhlik et al., 2009). Winter represents a significant energy restriction as deer have to move in the snow, face low temperatures, and cope with reduced forage quality and accessibility. In contrast, spring increases deer activity as they exploit the abundance of new vegetation and improved weather conditions (Bobek et al., 2016; Mysterud et al., 2001). Forage availability and distribution influence home range sizes due to climate change (Bobek et al., 2016; Van Beest et al., 2011), predator presence, or hunting season (Jarnemo & Wikenros, 2014). Other factors affecting home range size include age, sex, migratory strategies, supplemental feeding, and human disturbance (Börger et al., 2006; Luccarini et al., 2006; Mysterud et al., 2001; Reinecke et al., 2014). Red deer show significant variation in home range sizes across Europe (Jan F. Kamler et al., 2008), up to over 5000 ha seasonally (Náhlik et al., 2009), and to over 7000 ha annually (Zlatanova et al., 2019) depending on body mass due to higher metabolic needs (Makarieva et al., 2005; McNab, 1963). Spatial movement tends to be lower in resource-rich habitats, where nutritional gains are high, and travel costs are low (Börger et al., 2008).

Therefore, supplementary feeding can modify red deer spatial behavior, leading to smaller home ranges and changes in foraging habits and migrations (Felton et al., 2022; Jerina, 2012; Putman et al., 2004). Moreover, intensive feeding strategies mainly cause deer to establish home ranges near feeding sites, an effect intensified when natural resources are limited. This local aggregation increases pathogen transmission risks (Sorensen et al., 2014).

High deer densities lead to over-browsing, negatively affecting forest regeneration and biodiversity (Carpio et al., 2021; Čermák et al., 2004; Linnell et al., 2020; Valente et al., 2020). The effect of deer on woodland vegetation has resulted in significant changes in forest stand structure, composition, and stability (Gill & Beardall, 2001). The most known impact of red deer damage is browsing on young trees, which changes the stand structure and species composition and affects woodland development (Čermák et al., 2004; Cukor et al., 2022; Felton et al., 2022). Additionally, bark stripping is another severe damage that directly impacts the radial growth and productivity of affected forest stands (Cukor et al., 2019; Vacek et al., 2020).

Current advances in biologging technology and analytical methods provide an optimal solution for the precise recording of animal behaviors in the wild. These tools allow for non-invasive, long-term monitoring without the need for direct observations (Wilson et al., 2014). Biologging devices, such as GPS collars with accelerometers and magnetometers, enable continuous monitoring of animal movement and behavior, providing high-resolution data on spatial use and activity levels (Kirchner et al., 2023; Nathan et al., 2012; Williams et al., 2020).

In this study, we employ such technology to monitor the movement and energy expenditure of red deer, with a specific focus on the VeDBA (Vectorial Dynamic Body Acceleration) parameter (Wilson et al., 2019). Calculation works with raw acceleration excluding the gravitational component;

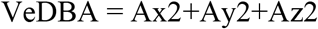

where A*x*, A*y*, A*z* are the dynamic components of each acceleration axis (Williams et al., 2020). This metric offers a detailed quantification of energy expenditure through the measurement of animal acceleration, providing a proxy for movement intensity and overall activity (Qasem et al., 2012; Wilson et al., 2014). Energy expenditure is a useful tool to measure the cost of specific activities that have a direct impact on fitness and natural selection, such as foraging, sleep, or movement (Gallagher et al., 2017; Qasem et al., 2012).

This study addresses a fundamental question: does the confinement of deer into seasonal enclosures reduce their movement and consequently their energy expenditure?

## Material and methods

### Study site and species

European red deer (*Cervus elaphus*), especially in winter and spring, exhibit a variety of spatial activities and energy requirements due to changes in environmental conditions and resource availability (Berger et al., 2002; Luccarini et al., 2006; Náhlik et al., 2009). Winter represents a significant energy restriction as deer have to move in the snow, face low temperatures and cope with reduced forage quality and accessibility. In opposition, spring increases deer activity as they exploit the abundance of new vegetation and improve weather conditions (Bobek et al., 2016; Mysterud et al., 2001). Forage availability and distribution influence home range sizes due to climate change (Bobek et al., 2016; Van Beest et al., 2011), predator presence, or hunting season (Jarnemo & Wikenros, 2014). Also, other factors affect home range size including age, sex, migratory strategies, supplemental feeding, and human disturbance (Börger et al., 2006; Luccarini et al., 2006; Mysterud et al., 2001; Reinecke et al., 2014). Red deer show significant variation in home range sizes across Europe (Jan F. Kamler et al., 2008), up to over 5000 ha seasonally (Náhlik et al., 2009), and to over 7000 ha annually (Zlatanova et al., 2019) depending on body mass due to higher metabolic needs (Makarieva et al., 2005; McNab, 1963). Spatial movement tends to be lower in resource-rich habitats, where nutritional gains are high, and travel costs are low (Börger et al., 2008).

Therefore, supplementary feeding can modify red deer spatial behaviour, leading to smaller home ranges and changes in foraging habits and migrations (Felton et al., 2022; Jerina, 2012; Putman et al., 2004). Moreover, intensive feeding strategies mainly cause deer to establish home ranges near feeding sites, an effect intensified when natural resources are limited. This local aggregation increases pathogen transmission risks (Sorensen et al., 2014).

High deer densities lead to over-browsing, negatively affecting forest regeneration and biodiversity (Carpio et al., 2021; Čermák et al., 2004; Linnell et al., 2020; Valente et al., 2020). The effect of deer on woodland vegetation has resulted in significant changes in forest stand structure, composition, and stability (Gill & Beardall, 2001). The most known impact of red deer damage is browsing on young trees, which changes the stand structure and species composition and affects woodland development (Čermák et al., 2004; Cukor et al., 2022; Felton et al., 2022). Also, bark striping is another severe damage that directly impacts the radial growth and productivity of affected forest stands (Cukor et al., 2019; Vacek et al., 2020).

Current advances in biologging technology and analytical methods provide an optimal solution for the precise recording of animal behaviours in the wild. These tools allow for non-invasive, long-term monitoring without the need for direct observations (Wilson et al., 2014). Biologging devices, such as GPS collars with accelerometers and magnetometers, enable continuous monitoring of animal movement and behaviour, providing high-resolution data on spatial use and activity levels (Kirchner et al., 2023; Nathan et al., 2012; Williams et al., 2020).

In this study, we employ such technology to monitor the movement and energy expenditure of red deer, with a specific focus on the VeDBA (Vectorial Dynamic Body Acceleration) parameter (Wilson et al., 2019). Calculation works with raw acceleration excluding gravitational component; 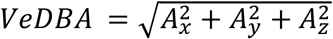 where Ax, Ay, Az are the dynamic components of each acceleration axis (Williams et al., 2020). This metric offers a detailed quantification of energy expenditure through the measurement of animal acceleration, providing a proxy for movement intensity and overall activity (Qasem et al., 2012; Wilson et al., 2014). Energy expenditure is a useful tool to measure the cost of specific activities that have a direct impact on fitness and natural selection, such as foraging, sleep or movement (Gallagher et al., 2017; Qasem et al., 2012).

This study addresses a fundamental question: does the confinement of deer into seasonal enclosures reduce their movement and consequently their energy expenditure?

## Material and methods

### Study site and species

The red deer (*Cervus elaphus*) enclosure management were conducted in the years 2019, 2020 and 2021 in the Czech Republic (Table 1). This study was realized in a state military training site which is closed for public and used only for military training, game species hunting and forestry. This location is situated at the Doupov and the area includes 331 km^2^. The red deer populations are managed by hunting for decades and the populations is stable. We chose 11 individuals in two seasons during the years 2019 and 2020 for this experiment and collected the spatial and movement data about. We used the winter enclosure (area 20 hectares) for this experiment. The winter enclosure is fenced area including pasture, bushes, forest stands and supplementary feeding sites inside. The enclosure is usually closed and managed by supplementary feeding since beginning of January until the end of April. The red deer individuals are able to jump inside the enclosure based on their decision during the managed season thru the one-way-entrance, but they are not able to go out until the managed opening time.

### Collaring

The immobilization for tagging were done by using the mixture of Ketamine and Xylazine then the individuals were tagged by GPS collar including the biologging sensor and battery. Animals handling was realized in accordance with the decision of the ethics committee of the Ministry of the Environment of the Czech Republic number MZP/2019/630/361 which approved the procedures.

### Data recording and tracking

We observed the red deer individuals by using GPS Vertex Plus collars (Vectronic Aerospace GmbH). The total weight of the collar was 750g and the lifetime was app. 1 year. The collar mass was lower than 3% of the animal’s body mass (Wilson 2021). The biologging data were recorded onboard SD cards with 16GB capacity. These data were downloaded after one year of observations when the animal was re-collared by the new GPS collar and new biologging tag. Once we had the collar back with one-year dataset we downloaded the biologging data. We also collected the GPS data in interval of 30 minutes during the observation period. The GPS data were updated remotely online on the server as a package of 8 hours-dataset.

Based on the GPS data we described the period when the individual was situated inside the winter enclosure and when the individual was released (direct management release) or escaped (not direct management release). We marked the time sequences inside the biologging data and used the information about spatial status as a factor for following analyses. In total we have data of 11 individuals and for 5 of them we were able to collect second season of tracking. The data of 8 individuals were recorded when the animals was inside the winter enclosure. The 3 individuals did not come inside the winter enclosure but were in close distance around. For all animals we have spatial and movement data from free range period. The 2 of 5 individuals were recorded two times inside the winter enclosure in the second year (Table 1). The winter enclosure was closed from 4/1/2019 until 29/4/2019 and from 28/1/2020 until 30/4/2020.

**Table 1.** Overview of collared red deer individuals.

| Animal ID | Time period | Data status | Sex | Year |
| --- | --- | --- | --- | --- |
| 107 | 01/04/2020 – 30/06/2020 | Inside | F | 2020 |
| 120 | 04/04/2020 – 30/06/2020 | Inside | F | 2020 |
| 117 | 12/03/2020 – 30/06/2020 | Inside | F | 2020 |
| 115 | 22/03/2020 – 07/04/2020 | Outside | F | 2020 |
| 108 | 28/03/2020 – 30/06/2020 | Inside | F | 2020 |
| 137 | 26/04/2020 – 30/06/2020 | Inside | F | 2020 |
| 135 | 20/03/2020 – 30/06/2020 | Inside | F | 2020 |
| 133 | 26/04/2020 – 30/06/2020 | Inside | F | 2020 |
| 104 | 21/04/2020 – 30/06/2020 | Outside | F | 2020 |
| 96 | 02/04/2020 – 30/06/2020 | Outside | F | 2020 |
| 151 | 30/03/2019 – 30/06/2019 | Outside | F | 2019 |
| 120 | 02/04/2019 – 30/06/2019 | Outside | F | 2019 |
| 117 | 27/03/2019 – 30/06/2019 | Outside | F | 2019 |
| 115 | 27/03/2019 – 30/06/2019 | Outside | F | 2019 |
| 108 | 31/03/2019 – 30/06/2019 | Inside | F | 2019 |
| 107 | 28/03/2019 – 30/06/2019 | Inside | F | 2019 |

### Data analyses

The biologging data were analyzed by daily summary of VeDBA values to evaluate the overall daily activity over 24 hours and by VeDBA sums in 60-minute intervals to assess activity in different parts of the day. The data were tested for normality, and differences were subsequently tested using one-way ANOVA followed by Tukey’s post-hoc test to determine differences between individual groups. A t-test was used to evaluate differences between two groups where applicable. The data were processed using Statistica, MS Excel, and R Studio. The ggPlot package was used for data visualization. For data comparison, we used the control period method (Bíl et al., 2018), where the control period was the time after the animals were released from the winter enclosure. Additionally, animals permanently living outside the winter enclosure were used as a control group.

## Results

### Total Daily Energy Expenditure

When comparing the total daily sum of VeDBA between animals that lived inside the winter enclosure and those outside (Fig. 1), a statistically significant difference was found only in March (t=6.181; p=0.000). During this period, the activity of animals in the winter enclosure was significantly lower compared to those living in the wild. The average VeDBA value for animals outside the winter enclosure was 4,473 ± 496, while for those inside the enclosure it was 107,615 ± 11,807. In April, the activity of both groups of animals equalized, and there was no statistical difference between them. The average VeDBA sum for animals in the wild in April was 114,679 ± 20,075, while for those in the enclosure it was 108,101 ± 35,905. However, the data variability for animals in the winter enclosure was very high, both in March and especially in April. The daily maximum VeDBA values for both groups in April were similar (13,341 and 13,703, respectively), but the minimum values for animals in the winter enclosure (43,238) were significantly lower compared to those living in the wild (min=93,042). The higher variability in the total daily VeDBA values for animals in the enclosure is also confirmed by the quartiles, which are almost twice as high as those for animals living in the wild (see Fig. 1). After the release of individuals from the winter enclosure, the daily energy sums were almost identical, and there were no differences between the two groups, with average daily VeDBA sums ranging from 94,414 ± 15,104 to 97,763 ± 13,353. When comparing the total daily VeDBA sum across individual months, April was statistically significantly different (36.893; p=0.000) from other months for animals living in the wild throughout the observation period, unlike animals from the winter enclosure, where their total daily activity differed statistically each month, with the highest again in April (F=20.415; p=0.000).

**Fig. 1.**
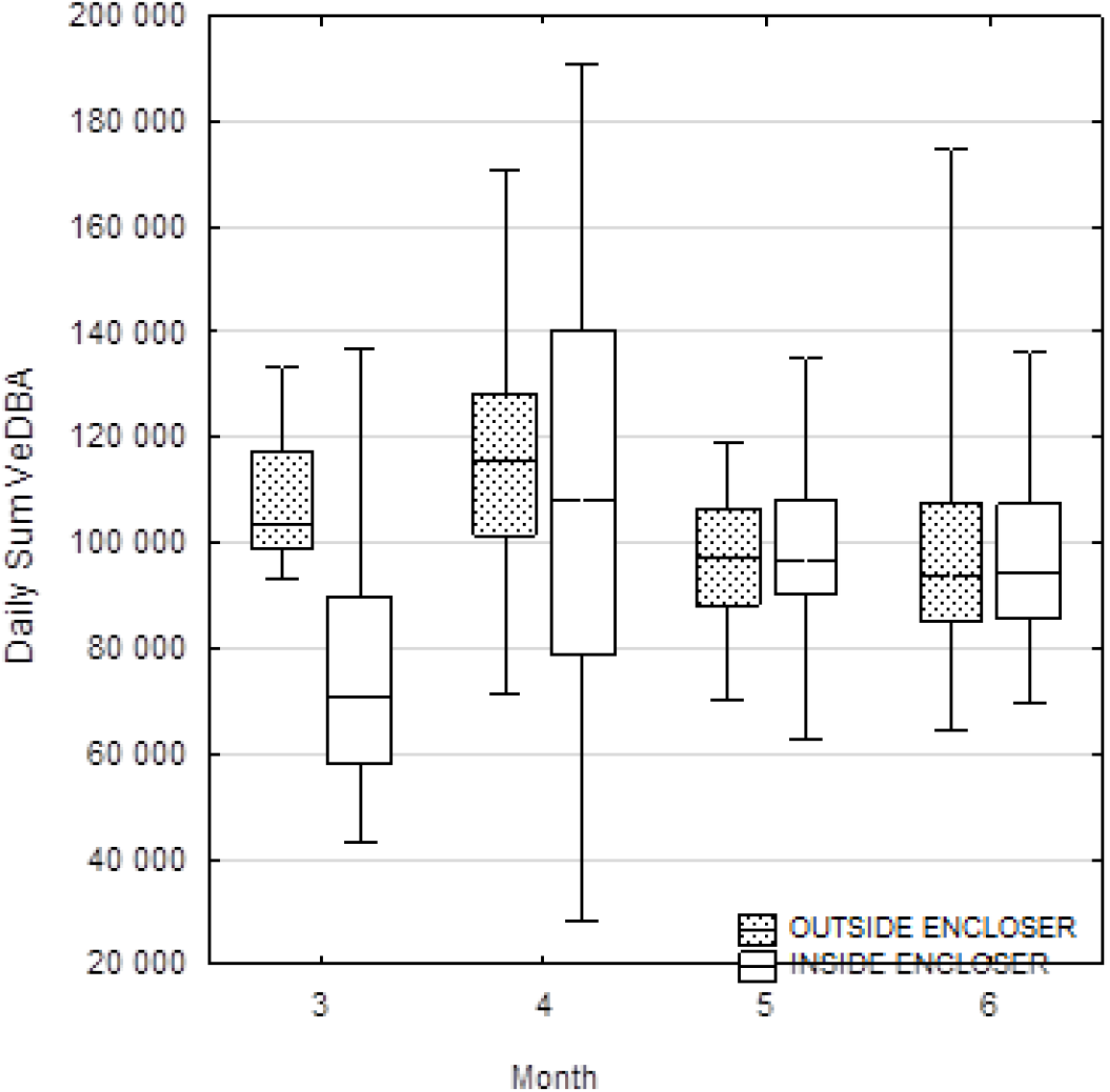
Daily sum of VeDBA for the animals which did not visit the winter encloser during March and April (OUTSIDE ENCLOSER) and for those who visited the encloser and were closed inside during March and April (INSIDE ENCLOSURE). Boxplots show the median values (middle bar in rectangles), upper and lower quartiles (length of rectangles), and maximum and minimum values (whiskers).

Some of the animals escaped from the winter enclosure during the period it was closed, and their activity significantly increased after leaving the enclosure, both in March and April (Fig. 2). This differs from the animals that remained inside and those that lived in the wild (March: F=44.415; p=0.000; April: F=4.897; p=0.007).

**Fig. 2.**
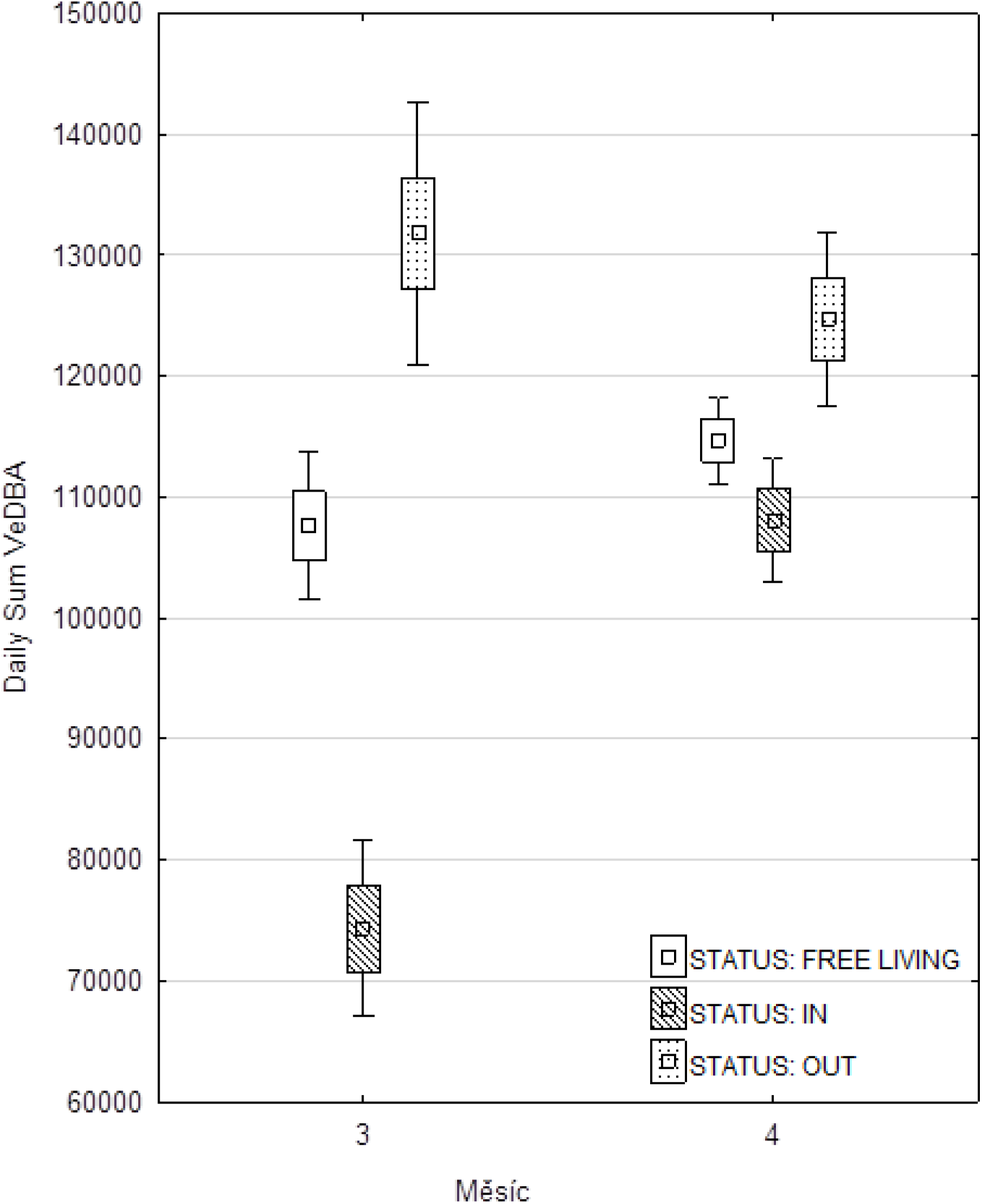
Daily sum of VeDBA for the animals which escaped from winter encloser during March and April (OUT); were closed inside (IN) and lived outside all the time). Boxplots show the mean values (middle bar in rectangles), standard error (length of rectangles), and 95% confidence intervals values (whiskers).

It appears that the period of release from the winter enclosure represents a significant moment for the animals. When evaluating the activity 15 days before and 15 days after the release, the activity 3 days before the release differed significantly from all other days (F=3.947; p=0.001). The VeDBA values increased significantly during these three days and returned to normal immediately after the release from the winter enclosure (Fig. 3).

**Fig. 3.**
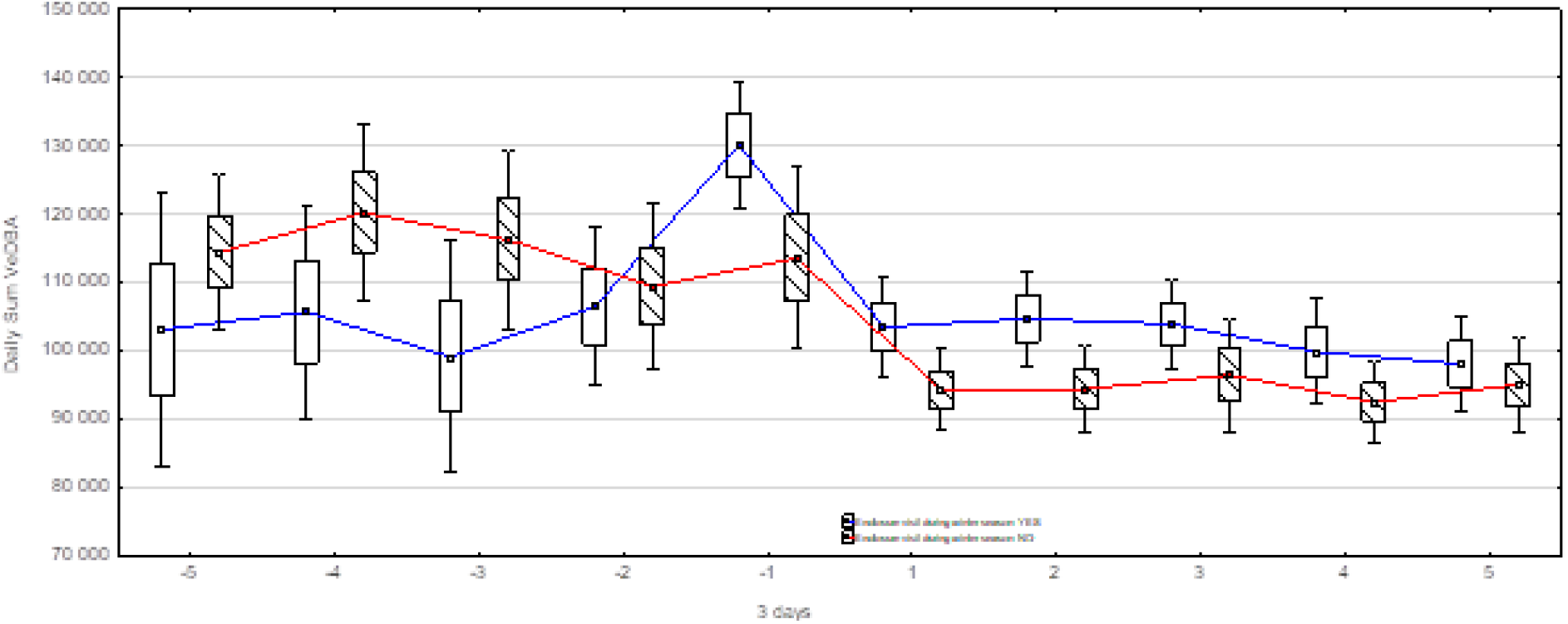
Daily sum of VeDBA for the animals 15 days before (-5 to -1, 1 step= 3 days) and 15 days after releasing from encloser (1 to 5) - red line and control group which were the same period outside (blue line) . Boxplots show the mean values (middle bar in rectangles), standard error (length of rectangles), and 95% confidence intervals values (whiskers)

### Daily Activity Pattern

The daily activity pattern of individuals in the winter enclosure shows no deviations from the daily activity pattern of deer living in the wild (Fig. 4). The peak activity occurs at sunrise and sunset. The lowest activity is shortly after midnight and then during the midday hours.

**Fig. 4.**
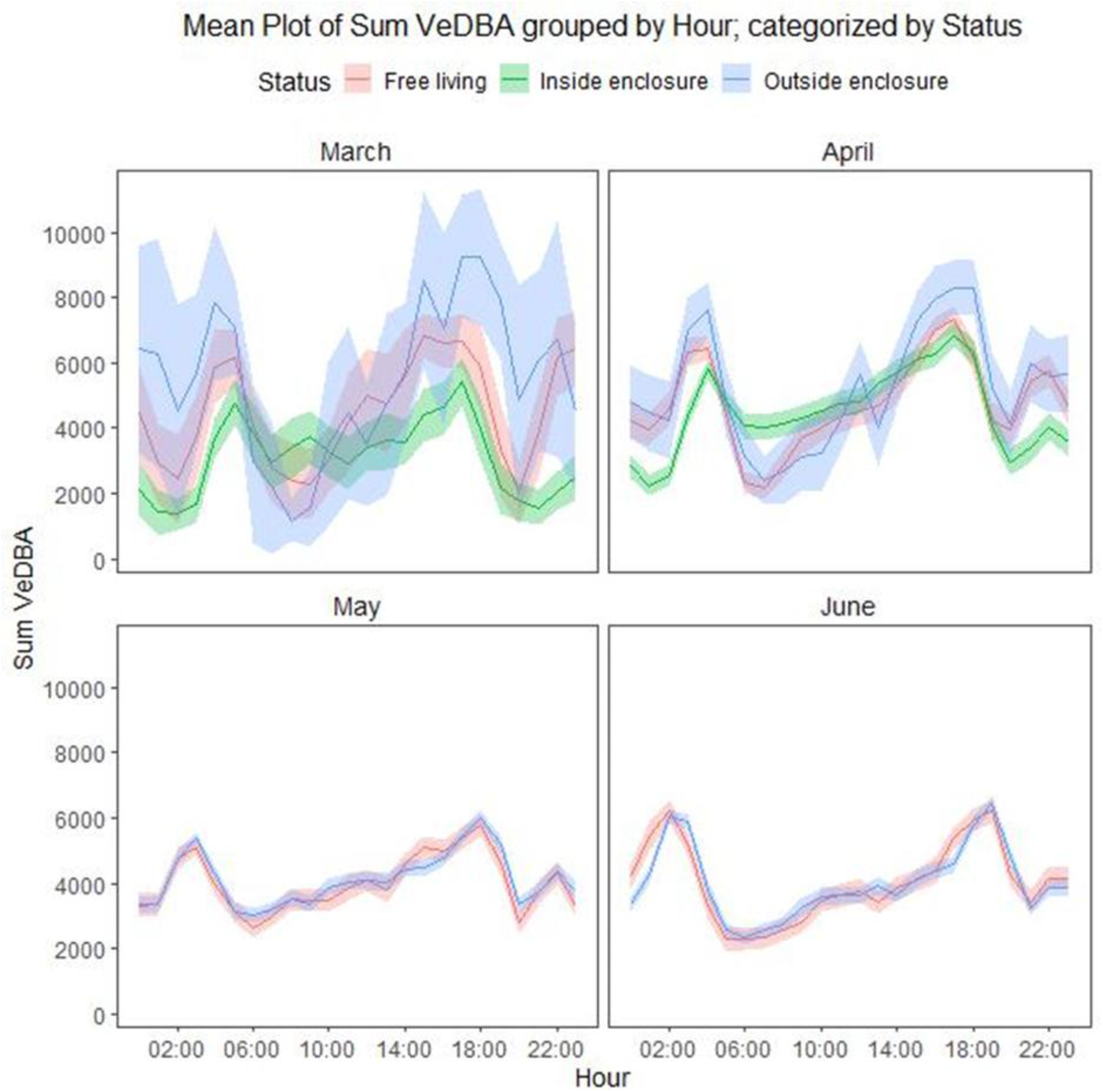
Hourly Sum of VeDBA during the day (24 hours) in different months.

Activity then gradually increases, reaching its peak during sunset. In March, it is evident that the highest activity is observed in individuals that left the winter enclosure, followed by those living in the wild, and the lowest activity is seen in individuals confined to the winter enclosure.

## Discussion

The activity of red deer throughout the year (seasons) and also during the day (24-hour cycle) has most often been evaluated using movement data, specifically based on the distance traveled during individual time intervals (Pépin et al., 2009; Georgii, 1980; Copes et al., 2017; Náhlik et al., 2009; Papon et al., 2013), or using activity sensors that measure activity over longer periods (Ensing et al., 2014; Lottker et al., 2009; Sunde et al., 2009). In recent years, high-speed accelerometers have been increasingly used to estimate energy expenditure – the so-called VeDBA (Goncalves et al., 2024; Benoit et al., 2020; Mortlock et al., 2024).

Estimating activity based on locomotion (i.e., distance traveled) is still the most common tool for evaluating activity. It is based on the assumption that locomotion causes a significant increase in energy expenditure and is much more energetically demanding than other types of behavior (Hudson, 1985). Therefore, most studies based on GPS telemetry report activity as the distance traveled between two GPS points. In this context, we hypothesized that individuals confined to the winter enclosure would have significantly lower energy expenditure than those living in the wild, primarily due to their limited movement. This hypothesis was confirmed for March but not for April. This likely relates to the increasing temperatures, which cause increased activity in cervids (Kamler et al., 2007). This is also confirmed by the research of Pépin et al. (2006), which showed increasing activity during spring and summer in captive red deer hinds, and by Reimoser et al., who confirmed increased heart activity from April to July, likely related to pregnancy and care for offspring. This suggests that increased energy expenditure is indeed dependent on general seasonal changes in the environment, and the confinement of wild animals in the winter enclosure does not significantly affect these processes.

We also expected that the daily activity pattern of animals confined to the winter enclosure would change significantly. Red deer living in the wild most often exhibit a circadian rhythm of activity, which is usually caused by anthropogenic pressure (Ensing et al., 2014; Vazquez et al., 2019; Hazlerigg and Tyler, 2019). This is confirmed by a study from the Białowieża Forest, where anthropogenic disturbance is minimal, and deer exhibited this rhythm only in winter, unlike the rest of the year (Kamler et al., 2007). In our study, the daily rhythm of individuals confined to the winter enclosure did not change and was consistent with the daily rhythm of deer in the wild. It is therefore evident that deer confined to the winter enclosure retain their daily cycles observed in the wild, and that confinement does not significantly reduce their wariness – i.e., their activity remains minimal during the day. The fact that the daily rhythm and behavior do not change significantly after confinement could be evidence of high welfare, indicating that wild animals do not suffer from being confined to the winter enclosure.

We observed a change in behavior only shortly before the individuals were released back into the wild. There was a significant increase in activity and thus energy expenditure in the last three days before release. This increase in activity just before release could be an indicator of how to maintain high welfare and could serve as a tool for determining when to release the animals back into the wild. According to a series of interviews with workers who care for animals in winter enclosures, the timing of release back into the wild is variable, and they usually recognize when the confined animals “want to go out” by observing changes in behavior, such as running around the fence and being significantly more restless than before.

It is therefore evident that their field observations and experience are based on real behavioral changes, which we confirmed. Therefore, long-term observation of individuals in winter enclosures is important for their welfare and maintaining natural rhythms as wild animals. The increase in activity could be one of the indicators of when it is necessary to release the animals.

## Conclusion

Our study demonstrated that red deer within winter enclosures exhibited significantly lower energy expenditure and reduced movement compared to those in free range, particularly in March. The average VeDBA values for deer inside the enclosures were substantially lower than those for free-ranging deer, indicating reduced activity levels. This suggests that the confinement within enclosures, combined with supplementary feeding, effectively limits the deer’s need to move extensively in search of food, thereby conserving their energy.

However, this difference in energy expenditure diminished in April as temperatures rose, suggesting that seasonal changes influence deer activity regardless of enclosure status. The increased temperatures likely lead to higher metabolic rates and greater activity as deer prepare for the more active spring and summer months.

Additionally, deer released from the enclosures showed a marked increase in activity and energy expenditure just before and after release. This spike in VeDBA values highlights the importance of timing the release to coincide with the availability of natural food resources. Proper timing ensures that the deer can transition smoothly back to free-ranging conditions without experiencing undue stress or energy deficits, which is crucial for their welfare and for minimizing potential damage to forest vegetation.

The use of winter enclosures, combined with supplementary feeding, effectively reduces deer movement and potential damage to young forest stands and seedlings during periods of low natural food availability. By concentrating deer in specific areas and providing consistent food sources, the enclosures help mitigate the negative impacts of over-browsing and bark stripping on forest regeneration.

Synchronizing the opening of enclosures with the availability of natural food resources is crucial for maintaining deer welfare and minimizing their impact on the environment. This approach not only supports the health and well-being of the deer but also protects forest ecosystems from excessive damage during critical periods of growth and recovery.

Overall, our findings suggest that winter enclosure management, when properly timed and supplemented with adequate feeding, can be an effective strategy for balancing the needs of wildlife conservation and forest management. Future research should continue to explore the long-term impacts of such management practices on both deer populations and forest health to refine and optimize these strategies further.

---

**Figure 7.** Female red deer with GPS collar.

**Figure 8.**
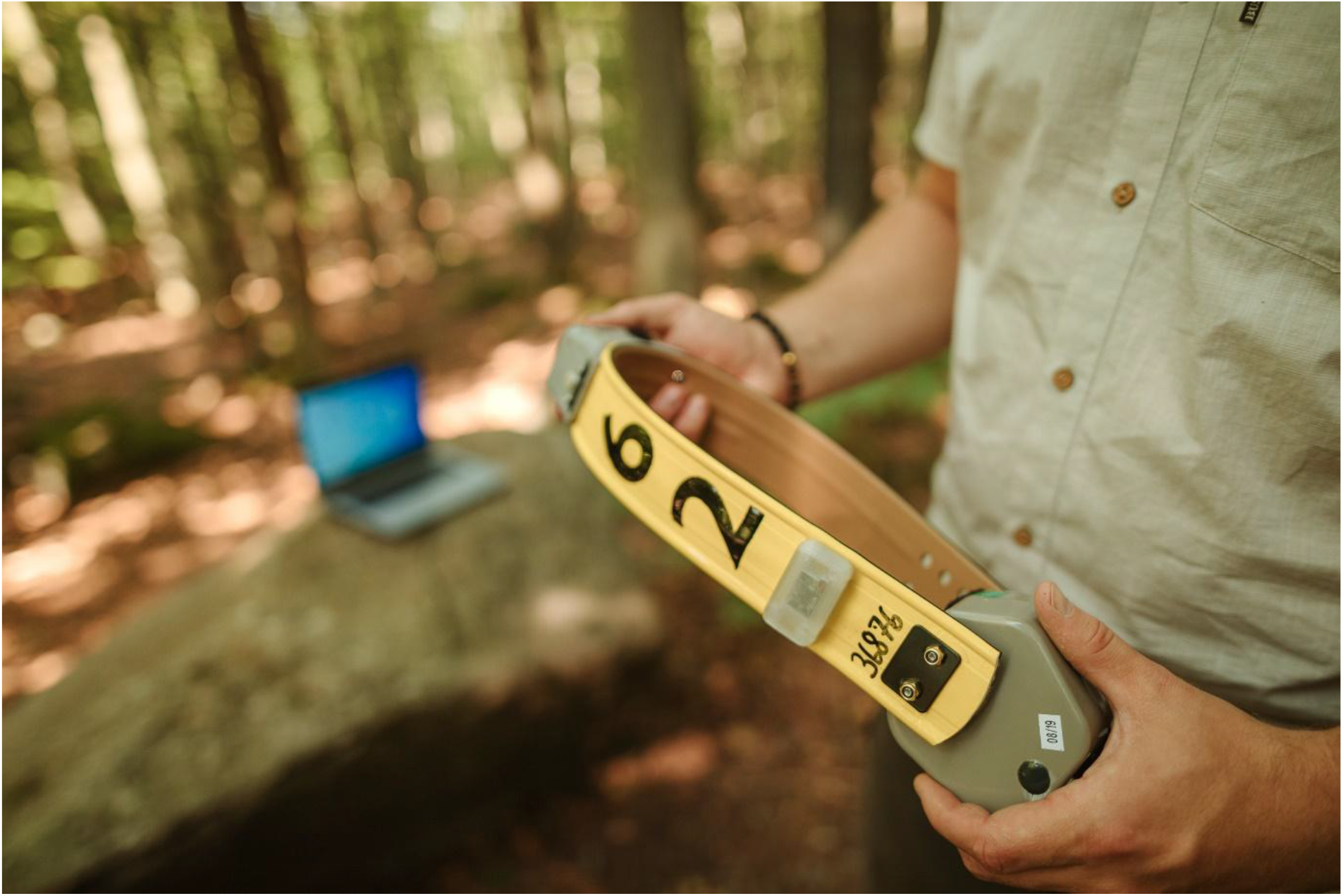
Detail of the used GPS collar including biologging sensor.

## Ethical approval

The red deer tagging was realized in accordance with the decision of the ethics committee of the Ministry of the Environment of the Czech Republic number MZP/2019/630/361.

## Acknowledgements

This study was supported by grant “EVA4.0” No. CZ.02.1.01/0.0/0.0/16_019/0000803 financed by OP RDE, grant No. QK1910462 financed by Ministry of Agriculture of the Czech Republic and Project B_19_02 of FFWS CZU. We are grateful to an anonymous reviewer for valuable comments and suggestions that improved the manuscript.

## Authors’ contributions

V.S. and M.J. originated the idea. V.S., M.F. and M.J. realized the trapping. V.S., M.F. and

A.P. wrote the manuscript with the contribution of M.J.

## Additional information

Supplementary information is available for this paper at **LINK**

Correspondence and requests for materials should be addressed to

### Figure legends

**Table 1.** Overview of collared red deer individuals

**Figure 1 –** Box plot of Daily mean VeDBA values of all individuals inside = 1 and outside= 0 of the enclosure

**Figure 2 –** Box plot of Daily Sum VeDBA values of all individuals inside = 1 and outside= 0 of the enclosure

**Figure 3 –** Box plot of Daily sum VeDBA values of all individuals inside = 1 and outside= 0 of the enclosure classified by months.

**Figure 4 –** Box plot of Daily mean VeDBA values of all individuals inside = 1 and outside= 0 of the enclosure classified by months.

**Figure 5 –** Box plot of Sum VeDBA values of all individuals inside = 1 and outside= 0 of the enclosure classified by hours.

**Figure 6 –** Box plot of Mean VeDBA values of all individuals inside = 1 and outside= 0 of the enclosure classified by hours.

**Figure 7.** Female red deer with GPS collar.

**Figure 8.** Detail of the used GPS collar

